# Flashzoi: An enhanced Borzoi for accelerated genomic analysis

**DOI:** 10.1101/2024.12.18.629121

**Authors:** Johannes C. Hingerl, Alexander Karollus, Julien Gagneur

## Abstract

Accurately predicting how DNA sequence drives gene regulation and how genetic variants alter gene expression is a central challenge in genomics. Borzoi, which models over ten thousand genomic assays including RNA-seq coverage from over half a megabase of sequence context alone promises to become an important foundation model in regulatory genomics, both for massively annotating variants and for further model development. However, its reliance on handcrafted, relative positional encodings within the transformer architecture limits its computational efficiency. Here we present Flashzoi, an enhanced Borzoi model that leverages rotary positional encodings and FlashAttention-2. This achieves over 3-fold faster training and inference and up to 2.4-fold reduced memory usage, while maintaining or improving accuracy in modeling various genomic assays including RNA-seq coverage, predicting variant effects, and enhancer-promoter linking. Flashzoi’s improved efficiency facilitates large-scale genomic analyses and opens avenues for exploring more complex regulatory mechanisms and modeling.

**Availability and implementation:** The Flashzoi model architecture is part of the MIT-licensed borzoi-pytorch package, can be found at https://github.com/johahi/borzoi-pytorch and installed via pip. Model weights for all four Flashzoi and Borzoi replicates are available at https://huggingface.co/johahi under the MIT license.

## Introduction

Modeling genomic readouts directly from DNA sequence is a powerful approach to investigate the genetic basis of gene regulation^1–3^. Deep learning models, in particular, have shown promise in elucidating the sequence determinants underpinning a variety of genomic assays including RNA-seq, DNase-seq, and ChIP-seq^2,3^. For mouse and human genomes, the availability of large compendia of genomic assays notably from the ENCODE^4^ consortium has enabled the training of models including Enformer^2^ and its recent successor Borzoi^3^. These models consider a sequence context of up to 500 kilobase pairs and jointly model thousands of coverage tracks of genomic assays. Since its release, Enformer has been widely used to predict effects of variants on enhancer and promoter activity, as well as on gene expression^5–8^. Moreover, Enformer and Borzoi have been employed as foundation models, i.e. as models upon which further models can be built for more specific tasks including personal gene expression^9^ as well as single-cell gene expression and accessibility^10–12^.

Enformer and Borzoi employ convolutional layers to capture local regulatory motifs. These condense the input into shorter representations (e.g., 128 bp tokens), which are then processed by transformer layers^13^. Transformer layers utilize self-attention to integrate long-range interactions between regulatory elements. A critical component of transformer layers is the incorporation of positional information, allowing the model to discern the relative positions of sequence elements. Both Enformer and Borzoi use relative positional encodings^14^. Borzoi uses a so-called central mask positional encoding which effectively results in a step function decaying away from 0. Nonetheless, the quadratic computational complexity of self-attention, where each part of the input sequence attends to every other part, limits the scalability of these models, especially for large datasets and computationally intensive tasks such as genome-wide variant effect prediction and the analysis of distal regulatory elements. The recently introduced FlashAttention-2 algorithm offers a way to faster and more memory-efficient transformer training and inference by improving usage of the fastest memory component of the GPU (SRAM) and computing attention in a tiled fashion^15^. However, FlashAttention-2 is incompatible with the handcrafted relative positional encodings used by Borzoi and Enformer.

Here, we introduce Flashzoi, a model that addresses these limitations. In Flashzoi, we replaced Borzois relative positional encoding with rotary positional encodings^16^, to leverage FlashAttention-2. We show that this change of the model increases training and inference speed by up to 3-fold while retaining or improving predictive performance.

## Methods

Flashzoi builds upon the U-net architecture of Borzoi^3^, a deep-learning model for predicting genomic readouts from DNA sequence. Borzoi processes 524 kilobases (kb) of DNA sequence using convolutional and max-pooling layers, resulting in 4,096 embeddings at 128 base-pair resolution, each with a dimensionality of 1,536. These embeddings are passed through 8 transformer blocks before being upsampled to 32-bp resolution using nearest-neighbor upsampling and convolutions. Separate output heads then generate predictions for human and mouse genomic assays (Figure 1a).

**Figure 1.**
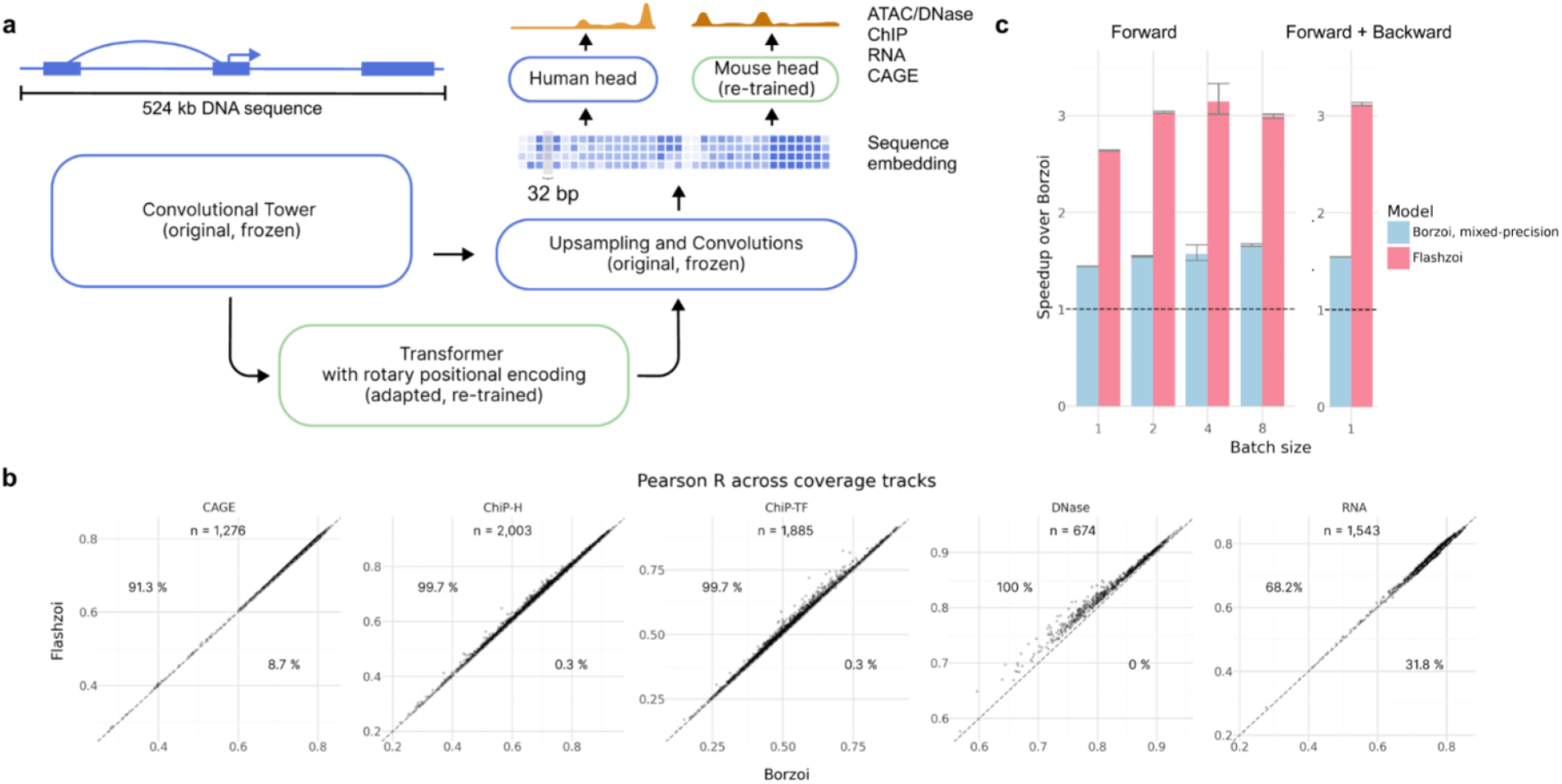
Flashzoi slightly improves genomic profile predictions over Borzoi at a 3-fold speedup. **a**, Flashzoi leverages architecture and pre-trained parameters from Borzoi, and replaces the Borzoi transformer with a FlashAttention-2 compatible one using rotary positional encodings. **b**, Pearson correlation between predicted and observed profiles on test sequences across genomic assays, for Borzoi and Flashzoi ensembles over four replicates. Percentages indicate proportions of tracks where Flashzoi outperforms Borzoi and vice-versa. Dashed line marks the y=x line. **c**, Speedup over Borzoi during forward, and forward and backward pass for a single Borzoi replicate run in mixed-precision, and Flashzoi. Dashed black lines indicate the Borzoi baseline (1.0), error bars indicate standard error (n=10).

The difference between Borzoi and Flashzoi lies in the transformer blocks. Each transformer block consists of a multi-head attention (MHA) module and a multi-layer perceptron (MLP). We modified the MHA module to use rotary positional encodings^16^, a widely used technique that allows attention between two tokens to depend on their relative distance. Our implementation uses FlashAttention-2.6.3^15^ and requires Nvidia GPUs of the Ampere generation or newer. We set the maximal frequency (theta) to 20,000 tokens and applied rotations to the first 128 dimensions of the query and key vectors per attention head. The remaining 64 dimensions were left unrotated which empirically improved performance.

While Borzoi’s MHA module projects queries and keys to 64 dimensions and values to 192 dimensions across 8 heads, Flashzoi uses grouped query attention^17^ (GQA) with 4 groups and 8 heads (192 dimensions each) to maintain a similar parameter count. This configuration allows one key and value head to be shared between two query heads, reducing the overall number of parameters without sacrificing model capacity. We also incorporated bias terms for the query, key, value, and output projections.

The training data for Flashzoi was downloaded from https://storage.googleapis.com/borzoi-paper/data/ and converted from the TFRecords format into WebDataset-compatible files for more efficient training. The human reference genome hg38 assembly, downloaded from the Borzoi repository^18^, was used for all analyses and training. We used gene annotations from GENCODE release v32^19^.

Borzoi comprises four independently trained replicates, each with a slightly different parameterization. For each Flashzoi replicate, we initialized all model parameters from the corresponding Borzoi replicate. Only the newly initialized transformer blocks and the mouse output head (absent in the original human-specific Borzoi weights) were trained. We adopted the same training-validation-test split (test fold 3, validation fold 4) as Linder et al.^3^ (test fold 3, validation fold: 4). We used the AdamW^20^ optimizer with a weight decay of 10^-8^ and a learning rate of 10^-4^, employing a global batch size of 8 across 8 GPUs, and froze the batch norms of all convolutional blocks. The model was trained using the same Poisson-Multinomial loss function as in Linder et al.^3^ with the same weighting factor (0.2), and batches alternating between human and mouse samples. Training was conducted using PyTorch^21^ 2.3.1 and FlashAttention^15^ 2.6.3 and stopped when the loss plateaued on the validation set. Training Flashzoi from scratch yielded inferior performance compared to initializing from Borzoi weights, a finding requiring further investigation in the future.

Borzoi was originally trained and evaluated in float32 precision. For consistency with the original publication, we also ran predictions for Borzoi in float32. However, we had to run Flashzoi in mixed-precision as the FlashAttention-2 floating-point format, which uses tensor cores, is half-precision (float16). Evaluations used FlashAttention 2.7.0 and PyTorch 2.5.1 (released after initial model training).

Genomic profile predictions were assessed using Pearson correlation between predicted and observed readouts on the held-out test set (fold 3). To quantify speedup, we measured the runtime of 10 forward and 10 forward-backward passes for single replicates of Flashzoi (mixed precision), Borzoi (float32), and Borzoi (mixed precision) across various batch sizes, with cuda.matmul.allow_tf32 enabled.

We evaluated the ability of Flashzoi and Borzoi to predict the effects of genetic variants on gene expression using fine-mapped GTEx eQTLs^22,23^. Only single nucleotide variants with a posterior inclusion probability greater than 0.9 were further considered. For each variant, the input sequence was centered on the TSS of the target gene (eGene) of the eQTL, and the predicted RNA-seq coverage track for the relevant tissue was obtained for both the reference and alternative alleles. Predicted variant effects were defined as the difference in logarithmic scale of the predicted gene expression of alternative and reference after removing the squashed scale^3^, summing across exon-overlapping bins, and adding a pseudocount of 1. The predicted variant effects were compared to observed variant effects (eQTL beta values) using Spearman correlation.

Furthermore, we assessed the performance of Flashzoi and Borzoi on eQTL prioritization using the tissue-specific datasets and matched negatives from Linder et al.^3^ Unlike their approach, which employed a random forest for Enformer comparison, we directly matched GTEx tracks for both Flashzoi and Borzoi. For this analysis, we followed the same variant-centered scoring strategy as Linder et al, in which a variant score was defined as the Euclidean norm across the 16,352 bins of the log_2_-fold change between the predicted RNA-seq coverage track for the alternative versus the reference after removing the squashed scale and adding a pseudocount of 1 on each bin.

Data for the Variant Flowfish^8^ analysis was downloaded from the supplementary material of the bioRxiv preprint. Indels were removed, and variant positions were lifted from hg19 to hg38. The effect of sequence perturbations on predicted RNA-seq coverage was recorded after removing the squashed-scale normalization for both Borzoi and Flashzoi.

To quantify the influence of distal regulatory elements on gene expression prediction, we utilized the CRISPRi benchmark dataset from Encode-E2G^7^, downloaded from https://github.com/EngreitzLab/CRISPR_comparison/blob/main/resources/crispr_data/EPCrisprBenchmark_ensemble_data_GRCh38.tsv.gz. This dataset provides a set of positive and negative enhancers for 1189 genes. For each enhancer-gene pair, the input sequence was centered on the transcription start site (TSS) of the linked gene. Predicted gene expression was calculated by summing the predicted readouts over all exons of the target gene. Following Gschwind et al., the gradient of the predicted gene expression with respect to the input sequence within 2 kb around the annotated enhancer midpoint was used as a proxy for enhancer importance^7^. Gradients were normalized by subtracting the mean nucleotide gradient. We calculated the area under the receiver operating characteristic (AUROC) for both Flashzoi and Borzoi when using the gradient score to predict positive enhancers.

## Results

We first evaluated whether Flashzoi maintained the predictive accuracy of Borzoi. Comparisons of genomic profile predictions on held-out sequences (test fold) revealed that an ensemble of four Flashzoi models always slightly yet consistently outperformed the four-model Borzoi ensemble across all data modalities (Figure 1b). Improvement also held when comparing a single instance of Flashzoi against a single instance of Borzoi. Most importantly, Flashzoi demonstrated up to a 3.2-fold speedup in training and inference time compared to Borzoi (Figure 1c). Moreover, Flashzoi exhibited reduced GPU memory consumption during both forward and backward passes (up to 1.4-fold for the forward pass, and 2.4-fold for forward and backward pass), indicating that Flashzoi could be used on servers with lower GPU-memory or with increased batch size. The performance gains are not solely attributable to Flashzoi’s use of mixed precision: even compared to Borzoi running in mixed precision, Flashzoi remains up to 2-fold faster during inference and backpropagation. These results demonstrate that Flashzoi achieves substantial performance gains without sacrificing predictive accuracy, making it a more efficient and practical tool for large-scale genomic analyses.

To assess generalizability and mitigate potential train-test set sequence similarity biases, we evaluated variant effect prediction using GTEx eQTLs. Across the 48 GTEx tissues in the eQTL catalogue^23^, Flashzoi predictions exhibited a significantly stronger Spearman correlation with observed eQTL effects compared to Borzoi predictions (*P* = 0.01, Figure 2a). We further assessed eQTL prioritization performance using distance-matched negatives from 49 GTEx tissues as previously collected by Linder et al. (2023). By ranking true eQTLs against matched negatives based on model predictions, we found the Flashzoi ensemble to be on par with the Borzoi ensemble in prioritizing eQTLs (Figure 2b).

**Figure 2.**
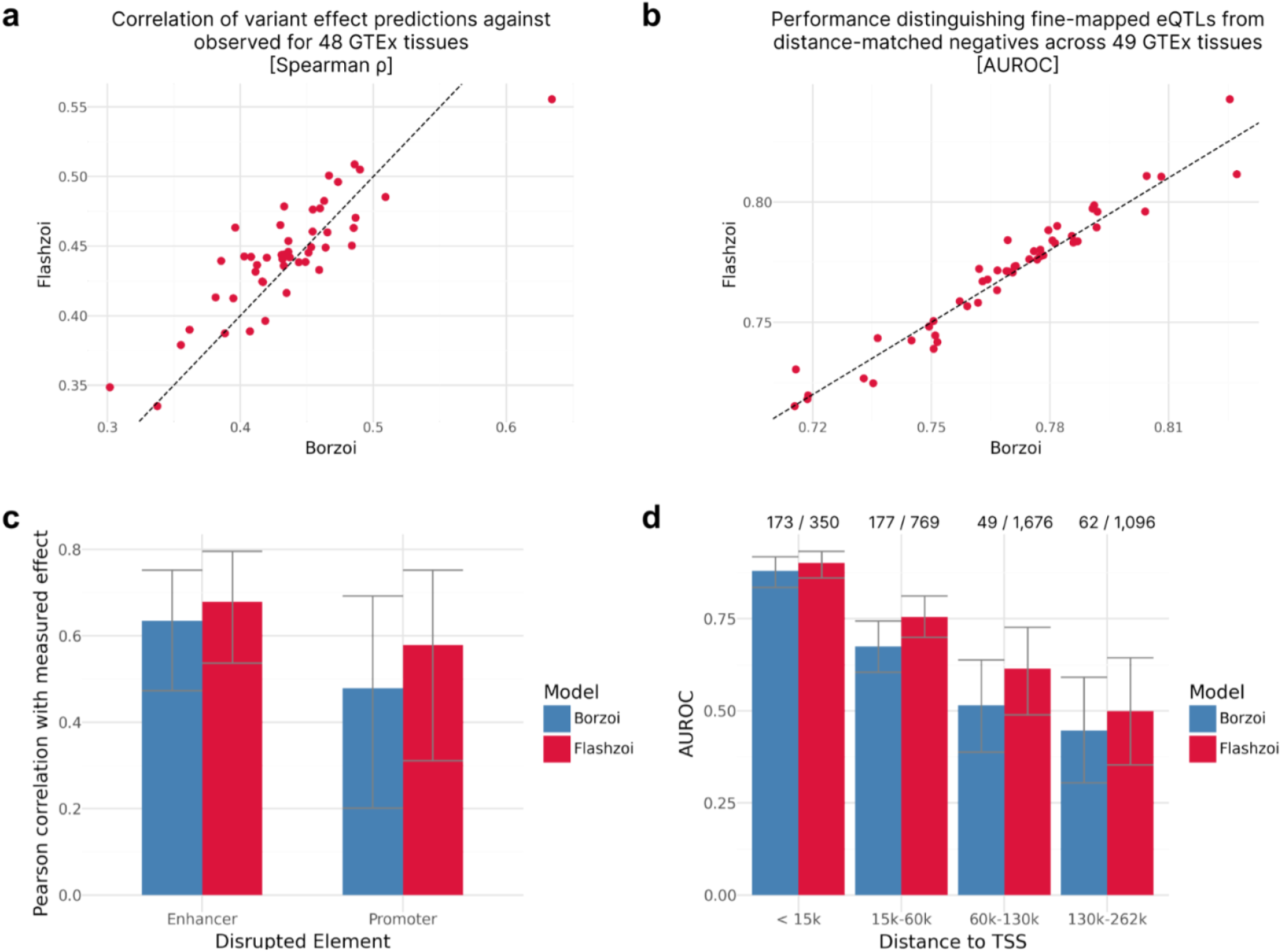
Flashzoi matches or outperforms Borzoi on variant effect prediction and better leverages distal elements for predictions. **a**, Spearman correlation of predicted effects (log-fold change) with observed GTEx normalized eQTL effects for Flashzoi against Borzoi. Each point indicates a GTEx tissue. Dashed line marks the y=x line. **b**, AUROC when distinguishing fine-mapped eQTLs from distance-matched negatives per GTEx tissue of Flashzoi and Borzoi. Dashed line marks the y=x line. **c**, Pearson correlation of Borzoi or Flashzoi prediction with measured effect of an enhancer or a promoter disruption on *PPIF* gene expression8. Error bars indicate 95% confidence intervals using bootstrapping. **d**, AUPRCs when using the gradients of Flashzoi or Borzoi to classify regulating CRE loci across genomic distances to TSS in data from Gschwind et al7. Error bars indicate 95% confidence intervals using bootstrapping. Proportion of significant CRE loci against the total number of CRE loci is indicated above each bar.

We also evaluated both models on two experimental datasets. Using the Variant Flowfish dataset^8^ which reports the regulatory impact of CRISPR-engineered variants in enhancers and promoters on expression of the *PPIF* gene, we found that Flashzoi predictions of *PPIF* were consistently on par with Borzoi for enhancer and promoter (Figure 2c). Finally, we analyzed the impact of CRISPR-inhibition of regulatory elements on gene expression using the ENCODE-E2G dataset^7^. Here too, Flashzoi was on par with Borzoi across all enhancer-promoter distances at distinguishing significant, experimentally confirmed enhancers from other enhancers (Figure 2d).

Altogether, these results show that Flashzoi maintains or slightly improves performance of Borzoi at much lower computational cost.

## Discussion

We introduced Flashzoi, an enhanced version of the Borzoi^3^ model for chromatin state and gene expression prediction. By leveraging rotary positional encodings and Flash Attention^15^, Flashzoi achieved an over 3-fold speedup and mildly improved predictive performance across various benchmarks. This substantial reduction in runtime and computational cost significantly benefits computationally intensive applications, including biobank-scale variant annotation, *in silico* sequence design, systematic investigation of regulatory elements, and the development of Borzoi-based foundation models.

We identified Borzoi’s transformer layers, specifically their positional encodings, as a computational bottleneck amenable to acceleration with FlashAttention-2. Further aspects could have been investigated. We considered using *compile*, the just-in-time compilation functionality of PyTorch, for various parts of the Borzoi model, but found it did not improve runtime. A promising future direction could be the use of Flex Attention^24^, a recently proposed compiler-driven programming model which allows exploring alternative attention variants. However, the current FlexAttention implementation does not support the exact dimensions of Borzoi.

Overall, Flashzoi is an efficient tool for accelerating research in gene regulation and variant interpretation, contributing to a deeper understanding of the noncoding genome. Flashzoi is a modification of Borzoi^3^. When using Flashzoi, we recommend citing the original Borzoi publication along with this manuscript.

## Acknowledgments

We thank Julia Maria Mayer for her cloud credits for downloading the Borzoi training data, and Laura Martens and Daniela Klaproth-Andrade for suggestions and feedback. We thank the authors of Borzoi for allowing us to upload PyTorch-ported Borzoi weights to the huggingface model hub. This study was funded by the European Research Council (ERC) (EPIC, Grant number: 101118521 to J.C.H. and J.G.) funded by the European Union. Views and opinions expressed are however those of the author(s) only and do not necessarily reflect those of the European Union or the European Research Council Executive Agency. Neither the European Union nor the granting authority can be held responsible for them. This study was funded by the Deutsche Forschungsgemeinschaft (DFG, German Research Foundation) via the project TRIUMPH - Tracking and Reconstructing Interactions to Understand the Missing Parts of Heritability (#535971044 to A.K.). This study was supported by the Deutsche Forschungsgemeinschaft (DFG, German Research Foundation) via the IT Infrastructure for Computational Molecular Medicine (Project #461264291).

## Notes

https://github.com/johahi/borzoi-pytorch

### Competing Interest Statement

The authors have declared no competing interest.

### Summary of Updates

Funding information was updated, and PPIF gene was spelled correctly.

